# Modulation of planarian seizure-like activity via antiseizure medications

**DOI:** 10.64898/2026.10.23.761305

**Authors:** Danielle Ireland, Eric Bormann, Evangeline Coffinas, Christina Rabeler, Eva-Maria S. Collins

**Author notes:** Corresponding author: (E-MSC).

## Abstract

Acute chemical exposure to organophosphorus compounds (OPs) arising from terrorist attacks or accidental poisonings pose a serious threat to human health. Acetylcholinesterase inhibition due to acute OP exposure causes a cholinergic crisis, manifested as increased secretions, respiratory distress, loss of consciousness, seizures, and ultimately death if left untreated. Benzodiazepines, such as midazolam, are currently used to stop OP-induced seizures, but must be administered within minutes after exposure to be effective. This makes midazolam impractical for large-scale civilian emergencies. Thus, there is a need to identify novel antiseizure medications (ASMs) that are equally effective even when administered later. It has been challenging to identify such compounds with traditional mammalian testing; high throughput screening assays that can quickly and cost-effectively evaluate many candidates for their potential as an ASM are needed. Planarians are aquatic invertebrates that are suitable organismal models for OP neurotoxicity studies due to the conservation of OP targets and their ability to metabolize OPs. Here, we built on our prior work that introduced a computational image analysis pipeline to quantify planarian seizure-like activity (pSLA) in two planarian species (*Dugesia japonica and Girardia dorotocephala)*. We previously used this approach to characterize pSLA following acute exposure (up to 30 minutes) to seizurogenic compounds. In the present study, we asked whether pSLA induced by the OP pesticide parathion, pentylenetetrazole, or NMDA could be rescued by co-exposure to four ASMs: midazolam, topiramate, MK-801, and carbamazepine. All tested ASMs were effective at reducing parathion-induced pSLA in one planarian species. However, ASM responses varied substantially across proconvulsants and planarian species. Defining the molecular mechanisms underlying pSLA will be essential for establishing planarians as a reliable model for ASM discovery.

## Introduction

Acute exposure to organophosphorus (OP) agents, such as nerve gases and OP pesticides (OPPs), resulting from terrorist attacks or industrial accidents represents a significant chemical threat and public health concern [1,2]. For example, a sarin attack on the Tokyo subway in 1995 injured over 5,000 people [3,4]. Acute OP toxicity results from inhibition of acetylcholinesterase (AChE), causing accumulation of acetylcholine and over-stimulation of nicotinic and muscarinic receptors. This cholinergic crisis manifests with increased secretions, respiratory distress, loss of consciousness, and seizures that evolve into status epilepticus, causing permanent brain damage and death if untreated [1,5]. Acute OP toxicity is currently treated with atropine and an oxime to mitigate cholinergic hyperstimulation [6,7], and a benzodiazepine, such as midazolam, to stop epileptic seizures [1,8]. However, midazolam is less effective in ameliorating seizures if administered after greater than 30-60 minutes of OP exposure [1,8] because of rapid internalization of synaptic GABA_A_ receptors following seizure onset, which progressively diminishes benzodiazepine efficacy [9,10]. Therefore, midazolam is impractical for large-scale civilian emergencies and there is a need for the development of novel antiseizure medications (ASMs) that are effective on longer time scales.

Development of new ASMs starts with *in silico* predictions of promising chemical structures, followed by *in vitro* tests of efficacy and toxicity, and subsequent mammalian testing of the most promising compounds [11]. These mammalian tests are slow and expensive, limiting the number of candidate compounds that can be tested [12]. Because *in vitro* systems cannot capture the metabolism and complexity of organisms, compounds that pass *in vitro* screening can still fail in animal tests, wasting time and resources. While these struggles are shared across drug sectors, identification of ASMs and therapeutics for acute OP toxicity also face the unique challenge that they cannot be ethically tested in human trials, making it very difficult to translate animal findings to clinical practice [13]. As a result of these limitations, despite intensive research into more effective therapies, the standard treatment for acute OP toxicity has remained the same for decades [13,14]. Thus, there is a need for organismal methods that are cost-effective and amenable to high-throughput screening (HTS) as an intermediate step to prioritize and filter promising ASM candidates [15–17]. Several organismal models have become popular for rapid screening of ASMs including fruit flies [18], nematodes [19] and developing zebrafish [12,20–23]. Of these, only developing zebrafish have been used to study seizure-like behavior following acute OP exposure [24,25], with limited tests of ASM efficacy [24]. Studies with fruit flies and nematodes often rely on genetic models and are better suited for epileptic research (reviewed in [17]). While developing zebrafish larvae are amenable to HTS, they do not reflect an adult nervous system and metabolism [12]. Thus, additional invertebrate organismal models that are suitable to HTS of seizure-like behavior in response to acute OP exposure in adult specimens are needed.

Like these other organismal models, freshwater planarians are amenable to low-cost rapid chemical screening [26–28]. While planarians have been used for decades to study how drugs affect brain regeneration, function, and behavior [29–33], they have recently become increasingly popular for both mechanistic studies [34–38] and medium-throughput screens of neuroefficacy and neurotoxicity of chemicals [26,27,39–43]. As an invertebrate, like fruit flies and nematodes, planarians can be studied both during development and in adulthood without ethical concerns or regulatory restrictions. With a few thousand neurons [44], the planarian nervous system is more complex than that of nematodes but simpler than that of developing zebrafish or fruit flies. Furthermore, *Dugesia japonica* (DJ) planarians are a mechanistically relevant model for studying OP neurotoxicity, because they possess two cholinesterase enzymes that are evolutionary ancestors of human AChE [36] and they metabolize OPPs similarly to humans; they are capable of OPP bioactivation and detoxification via carboxylesterase and paraoxonase activity [45]. Importantly, using multi-well behavioral screening and computational image analysis, we showed that the OPP parathion induced concentration-dependent planarian seizure-like activity (pSLA) in both DJ and *Girardia dorotocephala* (GD), another planarian species popular for chemical studies [40,41], within 30 minutes of exposure [46].

Here, we asked whether pSLA induced by parathion can be alleviated by midazolam, the currently approved ASM for acute OP exposure [8,14]. We also tested other ASMs that have previously been used in planarians: MK-801 [47], carbamazepine [48], and topiramate [47,49]. Prior manual studies in GD planarians found that NMDA and AMPA produce concentration-dependent pSLA, and that NMDA-induced pSLA was attenuated by the NMDA receptor antagonist MK-801, the AMPA receptor antagonist 6,7-Dinitroquinoxaline-2,3-dione disodium salt or the ASM topiramate [47]. Midazolam has been shown to produce dose-related physical dependence in GD planarians that is antagonized by the selective benzodiazepine receptor antagonist flumazenil [50], supporting the notion that functional GABA_A_/benzodiazepine receptor complexes are present and can be modulated pharmacologically. Moreover, it was found that carbamazepine also significantly reduced pSLA induced by either NMDA, nicotine or picrotoxin in *Girardia tigrina* (formerly *Dugesia tigrina*) planarians [48].

We hypothesized that pSLA induced by parathion or other seizurogenic compounds (pentylenetetrazole (PTZ), N-methyl-D-aspartate (NMDA)) can be rescued using one or more of these ASMs. We used the same computational analysis pipeline that we introduced recently [46] to quantify pSLA of both GD and DJ planarians and introduced new metrics to quantify effects on pSLA quantity and timing of onset. While our results demonstrate alleviating effects of some ASMs on pSLA intensity and/or onset, these effects were dependent on both the inducer and planarian species. Our quantitative approach enabled systematic evaluation of pharmacological modulation that was impossible with previous manual pSLA studies.

## Materials and methods

### Planarian species and maintenance

Experiments used the freshwater planarian species DJ or GD. DJ planarians have been maintained in the laboratory for > 15 years on a diet of organic liver (beef and chicken). GD planarians were purchased from Carolina Biological Supply (“Brown planarians”, Cat #132954, Burlington, NC, USA) and subsequently maintained on a diet of frozen bloodworms (San Francisco Bay Brand, Hayward, CA, USA, Cat #88055) in the laboratory. All species were maintained in planarian water (0.21 g/L Instant Ocean salts [Spectrum Brands, Blacksburg, VA, USA], 0.83 mM MgSO_4_, 0.9 mM CaCl_2_, 0.04 mM KHCO_3_, 0.9mM NaHCO_3_) [46]. Planarians were fed 1-2x per week and cleaned on the day of feeding and 2 days post-feeding. All experiments used intact, adult planarians (∼6 - 12 mm) which had been fasted 5-9 days prior to experiments. We have not observed differences in behavioral responses within this range of fasting state [46].

### Chemicals and preparation

Chemicals used in this study are listed in Table 1. Range finding experiments were first performed on the ASMs alone to determine concentrations that did not induce pSLA or other overt behavioral effects within 30 minutes of exposure in both species (S1-S2 Figures). These initial ranges (Midazolam: 0.316, 1, 3.16, 10, 31.6 µM; MK-801: 10, 31.6, 100, 316, 1000 µM; Topiramate: 31.6, 100, 316, 1000, 3160 µM; Carbamazepine: 3.16, 10, 31.6, 100, 316 µM) were guided by previous planarian literature on these compounds [26,47,48]. The maximum tested concentration with no behavioral effect alone was then used as the maximum concentration in co-exposures with the proconvulsants (Table 1). In co-exposure experiments, carbamazepine was tested in slightly different concentration ranges for the two planarian species due to differential sensitivity to carbamazepine alone (S1-S2 Figures). All other ASMs were tested in the same concentration range with the highest test concentration set as the highest concentration with no behavioral effects in both species. The proconvulsants (parathion, PTZ, and NMDA) were used at concentrations previously shown to produce robust pSLA in each planarian species within 30 minutes of exposure [46] (Table 1). For DJ planarians, parathion exposures were done with either 50 or 75 µM due to variability in potency of chemical stocks. These two concentrations induced similar levels and dynamics of pSLA and all data was normalized by experiment-specific parathion-only controls (see Statistical Analysis) to account for plate-to-plate variability. Depending on water solubility, chemical stock solutions were prepared either in planarian water or in 100% dimethyl sulfoxide (DMSO, Sigma-Aldrich, St. Louis, MO, USA) in a fume hood (Table 1). For all chemicals prepared in DMSO, the final DMSO concentration in all exposure solutions was 0.5% (v/v) and 0.5% (v/v) DMSO was used as the solvent control for these chemicals. DMSO at concentrations <4% do not induce pSLA in these planarian species [46]. Chemicals prepared in planarian water were used the same day. Stocks made in DMSO were aliquoted and stored at −20℃ or −80℃ until use. Frozen stocks were thawed a maximum of 3 times [51].

**Table 1.** Overview of chemicals and concentrations used.

| Chemical | CAS # | Tested concentrations<sup>a</sup> | Vendor | Purity (%) |
| --- | --- | --- | --- | --- |
| Carbamazepine (CAR)<sup>b</sup> | 298-46-4 | 10, 31.6, 100, 316 µM (DJ)<br>3.16, 10, 31.6, 100 µM (GD) | Cayman Chemical | ≥98% |
| Midazolam (MDZ)<sup>b</sup> | 59467-70-8 | 0.1, 0.316, 1, 3.16 µM | Sigma-Aldrich | 100% |
| (+) MK-801 hydrogen maleate (MK)<sup>b</sup> | 77086-22-7 | 1, 3.16, 10, 31.6 µM | Cayman Chemical | ≥98% |
| Topiramate (TOP)<sup>c</sup> | 97240-79-4 | 100, 316, 1000, 3160 µM | Cayman Chemical | ≥98% |
| N-methyl-D-aspartate (NMDA)<sup>c</sup> | 6384-92-5 | 3 mM (GD);<br>15 mM (DJ) | Tocris Bioscience | ≥99% (HPLC) |
| Parathion (PT)<sup>b</sup> | 56-38-2 | 31.6 µM (GD);<br>50, 75 µM (DJ) | Sigma-Aldrich;<br>ChemService | ≥98.0% (HPLC);<br>98.8% (GC/FID) |
| Pentylentetrazole (PTZ)<sup>c</sup> | 54-95-5 | 15 mM | Cayman Chemical | ≥98% |

<sup>a</sup> If the same concentration or concentration range was used in both species, there is no species reference; else, it is indicated in parentheses which species was tested at this concentration.
<sup>b</sup> Prepared in DMSO.
<sup>c</sup> Prepared in planarian water.

We measured the pH of the highest tested concentration of each chemical alone and in the co-exposure mixtures using an Apera PH60-MS pH Tester kit (Apera Instruments, Columbus, OH, USA) after allowing the probe to stabilize >10 minutes. The pH of the highest tested concentration of each chemical alone and in the co-exposure mixtures are listed in S1 and S2 Tables, respectively. NMDA, alone and in combination, at the concentrations used here (3 and 15 mM for GD and DJ, respectively) showed very low pH values (2.35-3.57) which we have previously shown can induce pSLA alone and likely contributes to NMDA-induced pSLA [46]. All other chemicals were within an acceptable pH range (4.83-8.10) that we have previously shown does not induce acute behavioral effects in DJ planarians [26]. Some of the measured concentrations are higher than ultimately used in this study, but since no obvious effects on pH were seen, the pH of lower concentrations were not recorded.

### Exposure and behavioral recordings

Individual planarians were each loaded into a well of a tissue culture treated 48-well plate (Genesee Scientific, El Cajon, CA) in 100 µL planarian water. 100 µL of 2x chemical stocks, prepared immediately before loading planarians into the 48-well plate, were added to bring each well to 200 µL at 1x the final concentration. Each plate tested 1 planarian species across 6 conditions, 1 condition per row (n=8 per plate). The conditions consisted of vehicle control (VC), proconvulsant alone, and 4 combinations of proconvulsant/ASM across different concentrations of ASM (Fig 1A). If the inducer and/or ASM were prepared in DMSO, the VC was 0.5% (v/v) DMSO, else it was untreated planarian water. Chemical addition was performed in the fume hood using a multichannel pipettor (CAPP Aero 12-Channel Pipette, 30-300 µL). After chemical addition, the plate was sealed using thermal film (Excel Scientific, Victorville, CA, USA). Three independent replicates (separate plates made from independent chemical stocks) were tested for each condition and species, with the relative orientation of the conditions across rows shifted between replicates to control for positional effect, for a total n=24. We have previously shown that this sample size has sufficient statistical power to differentiate pSLA occurrence [46]. Some NMDA-containing conditions were run in additional plates to increase sample size due to a high propensity of NMDA-exposed planarians to crawl out of their wells within 5 minutes of exposure.

**Fig 1.**
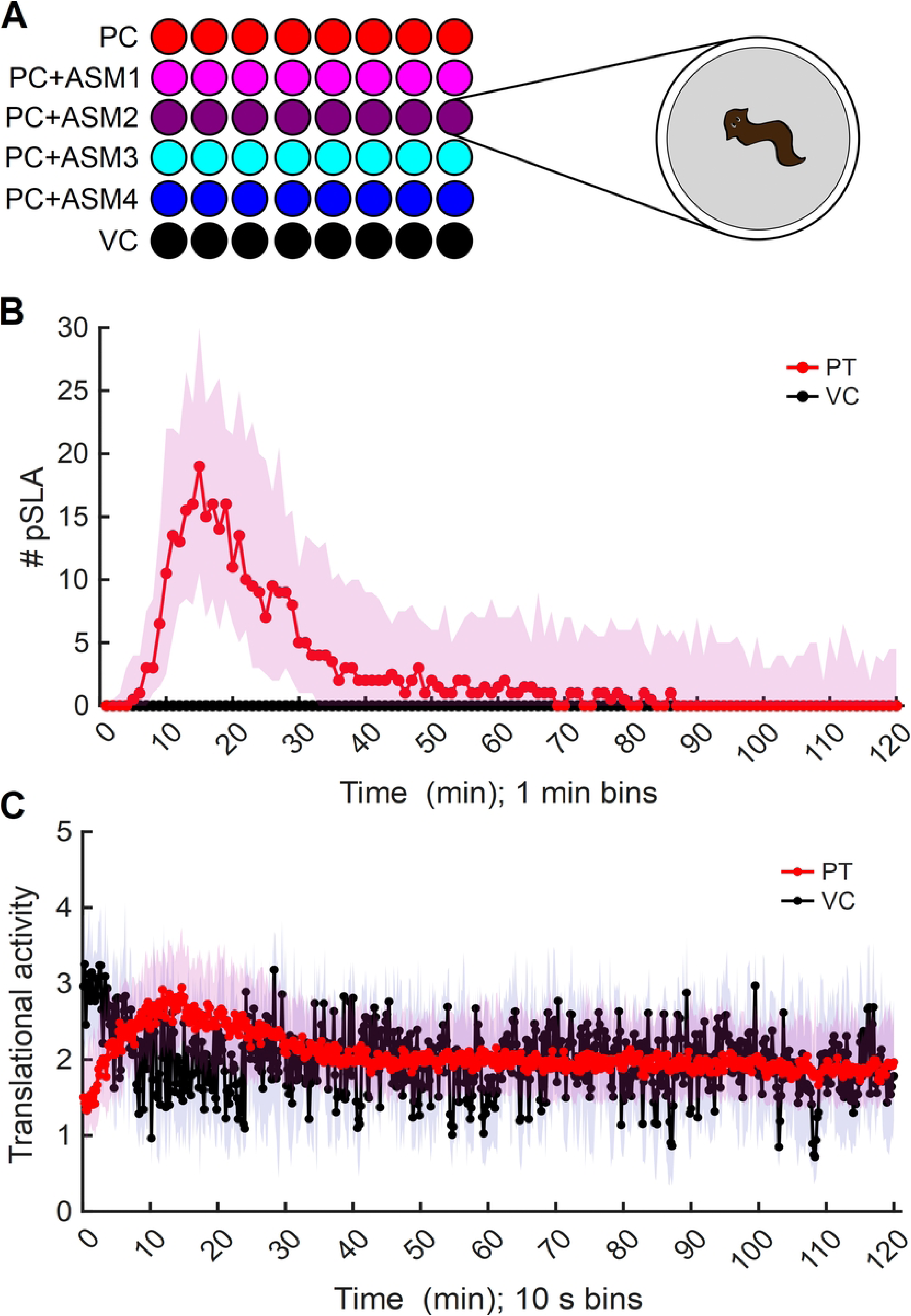
Plate overview and example parathion behavior. A) Schematic of conditions in each 48-well plate. Each row tested n=8 planarians in the same condition (1 worm per well). PC: proconvulsant (Parathion, PTZ, or NMDA); ASM: antiseizure medication (midazolam, topiramate, MK-801, carbamazepine; each tested at 4 concentrations in half-log steps, see Table 1 for concentrations). VC: vehicle control (planarian water or 0.5% (v/v) DMSO). B-C) Progression of behavioral phenotypes following exposure to parathion for 2 hours in DJ planarians, quantified via pSLA (B) and translational activity (C). Data points indicate the median and shading indicate the interquartile range from data compiled across two independent experiments (N_PT_=68; N_VC_=16).

Behavioral recordings were primarily taken 2-3 minutes after chemical addition (2 of the dose finding tests were taken within 5 minutes) with a custom imaging setup inside a NEEWER Photo Studio Shooting tent (Amazon.com, Seattle WA) as previously described [46]. Plates were set on an LED light panel with brightness set to 1350 to 1450 lux, as measured with a lux meter, and a Fresnel lens was placed on top of the plate to minimize imaging aberrations. Images were collected at 5 frames per second (fps) using a Point Grey Flea3 camera (FLIR, Wilsonville, OR) using Flycap software. Experiments with NMDA were recorded for 5 minutes and with PTZ for 20 minutes. Initial dose range finding experiments and experiments with parathion were 30 minutes long. These durations were tuned to each inducer to allow for the minimum amount of time to observe robust pSLA in both species and avoid confounding factors, such as crawl-outs in the case of NMDA [46]. For example, in DJ planarians exposure to parathion induces robust pSLA within 5-20 minutes, which dissipates by approximately 40 minutes (Fig 1B), replaced by less translational motion as the worms slowly twitch in place (Fig 1C).

### Automated image analysis

Analysis of the behavioral recordings was performed in MATLAB (2025a, MathWorks, Natick, MA) using custom scripts as previously described [46], with some modifications. Any worm that crawled out of the water during recording was excluded from analysis. A pSLA event was quantitatively identified as an instance where the median normalized motility index (MI) score for a 1-second (5-frame) bin exceeded a previously determined species-specific threshold [46]. The number of pSLA events across the entire recording period was summed. To quantify the rate of pSLA induction in DJ planarians exposed to parathion or PTZ, the slope of a linear trendline against the number of pSLA events per minute (ΔpSLA/min) during the time period when the number of pSLA per minute generally increases linearly (Fig 1B) was calculated for each planarian. We found that the timing of pSLA onset could vary from plate to plate. Therefore, the exact time range of interest was automatically chosen for each plate using findchangepts, using the linear statistic, in MATLAB on smoothed data of the median number of pSLA per minute of the proconvulsant alone. The time points were manually verified and updated if necessary. Translational activity was also quantified for each planarian over 10-second (50 frame) bins and normalized by worm area as previously described [46] (Fig 1C). The bin sizing was decreased from 20 to 10 seconds here to better ensure that gliding planarians did not glide back over their tracks, which would underestimate translational activity. In plots of translational activity over time, the median across all experiments is shown. For each planarian, the area under the curve (AUC) of the translational activity over time was quantified using the trapz function in MATLAB.

### Statistical analysis

Statistical analysis was performed in R (4.5.0, [52]). Sample sizes and p-values for all experiments are listed in S3 Table. Raw and analyzed data have been deposited on zenodo.org (doi: 10.5281/zenodo.23480117).

#### Range-finding tests

The number of pSLA events over the entire recording was counted and the data across all replicates was compiled. To determine if ASM concentration had a statistical effect on the number of pSLA events, a negative binomial generalized linear model was performed using the MASS package [53]. Post hoc pairwise comparisons of the estimated marginal means of the model was performed between the VC and each ASM concentration using the contrast function of emmeans [54]. The Benjamini & Hochberg (BH) adjustment method was used to adjust the p-values for multiple comparisons.

#### Inducer/ASM co-exposures

The number of pSLA events over the entire recording period was counted. Because the number of pSLA events induced by the same proconvulsant could differ considerably between replicates, the number of pSLA events within each plate was normalized by dividing by the median number of pSLA events of the proconvulsant alone before combining the data from all replicates. Translational AUC scores were also normalized by the median values of proconvulsant alone from the same plate before combining data across replicates. For statistical comparisons of normalized pSLA counts, ΔpSLA/min, and translational AUC scores, a Kruskal Wallis omnibus test, followed by a post hoc one-sided (effects in the decreasing direction) Dunn’s test comparing each condition to the proconvulsant alone was performed using the PMCMRplus package [55]. The BH adjustment method was used to adjust the p-values for multiple comparisons.

## Results

### Midazolam attenuation of parathion-induced pSLA is species-specific

Range finding tests showed that midazolam at concentrations of 10 µM and higher induced pSLA in DJ planarians (S1 Fig). Co-exposure of lower concentrations of midazolam (0.1-3.16 µM) with parathion significantly decreased the number of pSLA events compared to parathion alone (Fig 2A). To gain more nuanced insight into the dynamics of pSLA induction, we also studied the number of pSLA events per minute over the course of the 30-minute recording (Fig 2B) and calculated the slope of pSLA events per minute (ΔpSLA/min) during the linearly increasing portion of the response, with the exact timing tuned to the in-plate parathion only controls (Materials and Methods) (Fig 2C). These data showed that co-exposure to all concentrations of midazolam had significantly reduced induction of pSLA compared to parathion alone (Fig 2B-C). As we noticed that pSLA seemed to be increasing in some of the co-exposures at the end of the 30-minute recording, we also tested parathion and midazolam co-exposure with recording for 90 minutes (S3 Fig). These longer recordings revealed that while co-exposure to parathion and 0.1 µM midazolam never significantly induced pSLA, higher concentrations of midazolam only delayed parathion-induced pSLA, and pSLA in the co-exposure conditions eventually reached the same levels as with parathion alone. Thus, these 90-minute recordings provide insight into the dynamics of midazolam efficacy, but they do not alter the interpretation of the 30-minute exposures: either a loss or delay in pSLA induction would be considered a therapeutic effect. Therefore, 30-minute recordings were deemed sufficient to identify potential therapeutics for parathion-induced pSLA and were used for all subsequent experiments.

**Fig 2.**
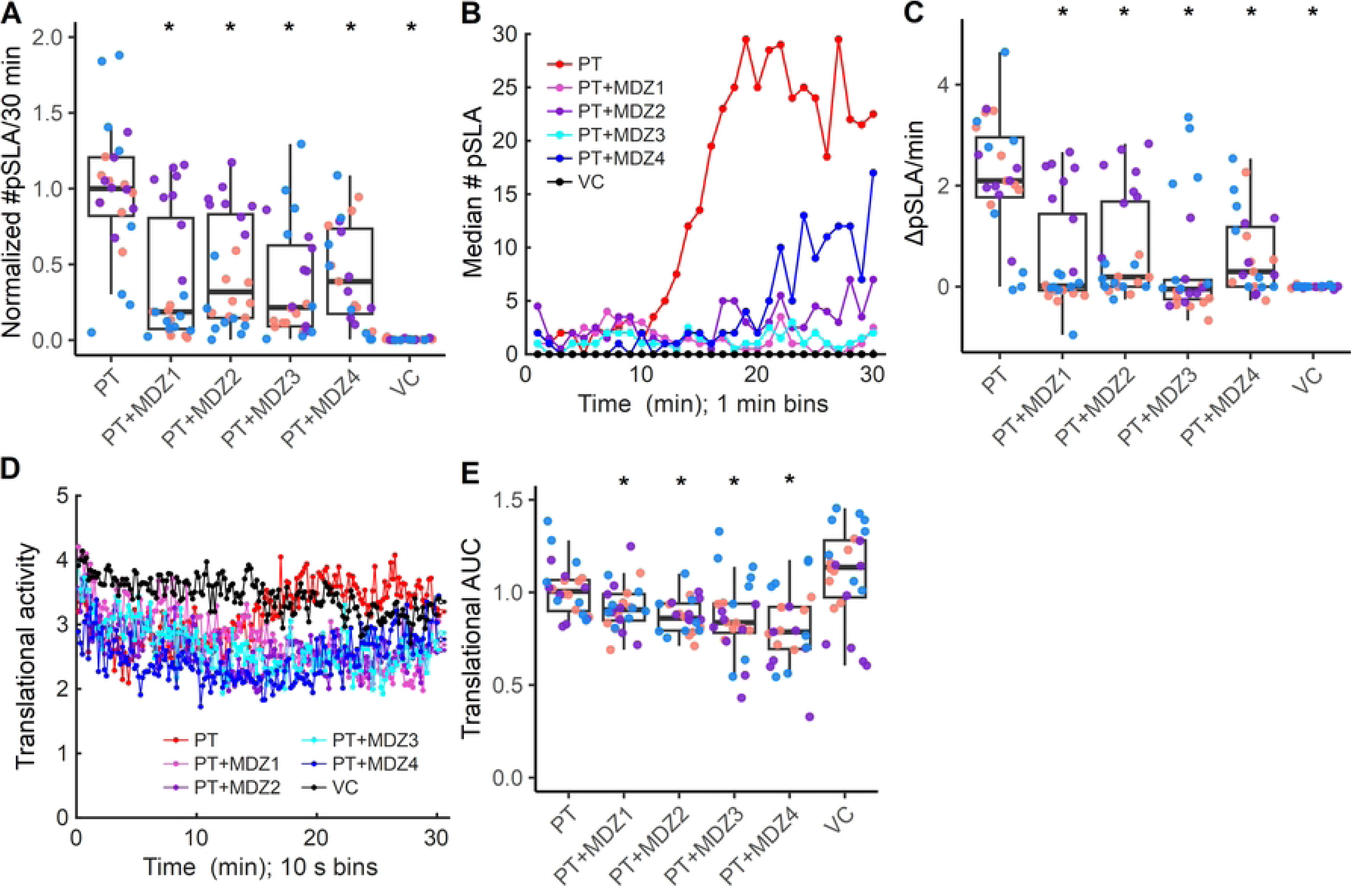
Midazolam decreases parathion-induced pSLA in DJ. A) DJ planarians were co-exposed to 50 or 75 µM parathion (PT) and varying concentrations of MDZ (MDZ1-4 correspond to 0.1, 0.316, 1, and 3.16 µM, respectively) and the total number of pSLA events, normalized by in-plate parathion-only controls over the 30-minute recording were counted. VC: vehicle control. B-C) Dynamics of pSLA induction were visualized by (B) plotting the median number of pSLA for each minute and calculating (C) the ΔpSLA/min during the linearly increasing time period, determined by the parathion-only controls from each plate. D-E) Translational activity was quantified from minimum intensity projections and normalized by each worm’s area. Plots of translational activity over time (D) show MDZ co-exposures decrease their translational activity over time and is quantified by comparing the area under the curve (AUC) (E). Asterisk (*) indicates statistical significance (p<0.05) using a Kruskal Wallis omnibus test, followed by a post hoc one-sided Dunn’s test comparing each condition to the inducer alone with p-value correction using the BH method. Color-coding of individual data points in the boxplots indicates indicates 3 independent experiments.

Because loss of pSLA with ASM co-exposure could be due to true therapeutic effects or due to high toxicity leading to loss of all movement, we also quantified effects of ASM co-exposure on translational activity to differentiate these phenotypes. Co-exposure with all concentrations of midazolam significantly decreased translational activity compared to parathion alone (Fig 2D-E). Focusing on co-exposure with 0.1 µM midazolam, we observed that this decrease in translational motion was due to the worms generally moving less and more slowly, but without any obvious effects on morphology to indicate high toxicity (S1 Movie). Because midazolam is a sedative that is expected to reduce locomotor activity [56], the observed decrease in translational motion is consistent with its known pharmacological effects rather than being indicative of overt toxicity.

In contrast, in GD planarians, co-exposure to 0.1-3.16 µM midazolam had no significant effects on parathion-induced pSLA or on translational activity compared to parathion alone (S4 Fig). It was not possible to evaluate effects on pSLA dynamics with GD planarians as they immediately began performing pSLA, which quickly dissipated as the planarians transitioned to intermittent gliding, scrunching, and other abnormal behaviors before eventually becoming contracted (S4B Fig), also leading to overall fewer pSLA events than with DJ planarians exposed to parathion (median in DJ: 472 vs median in GD: 22). The lower numbers of pSLA in GD planarians also led to greater variability across and within experiments. Notably, midazolam alone also had a weaker effect in GD planarians. Unlike in DJ planarians, midazolam concentrations up to 31.6 µM did not induce pSLA in GD planarians (S2A Fig). However, 31.6 µM midazolam reduced GD translational motion, suggesting that the lack of effect was not due to a lack of bioavailability.

### Topiramate, MK-801 and carbamazepine modulate parathion-induced pSLA in different ways

We also tested whether other ASMs which have been previously shown to be effective at mitigating pSLA induced by NMDA in some planarian species could alleviate parathion-induced pSLA (Fig 3). Co-exposure of parathion and either 1000 or 3160 µM topiramate significantly decreased the number of pSLA events in DJ planarians (Fig 3A). Additionally, 100, 1000, and 3160 µM topiramate also decreased the rate (ΔpSLA/min) of pSLA induction. While the effects of co-exposure with 100 µM topiramate were not clear in the time-dependent plot (Fig 3Aii) and the slope showed large variability among individuals, the slope was significantly reduced compared to parathion alone (Fig 3Aiii). Co-exposure with 3160 µM topiramate significantly decreased translational activity (S5A Fig) as the planarians became rigid/stationary, with intermittent twitching, abnormal morphology and pharynx extrusion (S5D Fig), suggesting toxicity. Together, these data show that 1000 µM topiramate causes a delay in parathion-induced pSLA in DJ.

**Fig 3.**
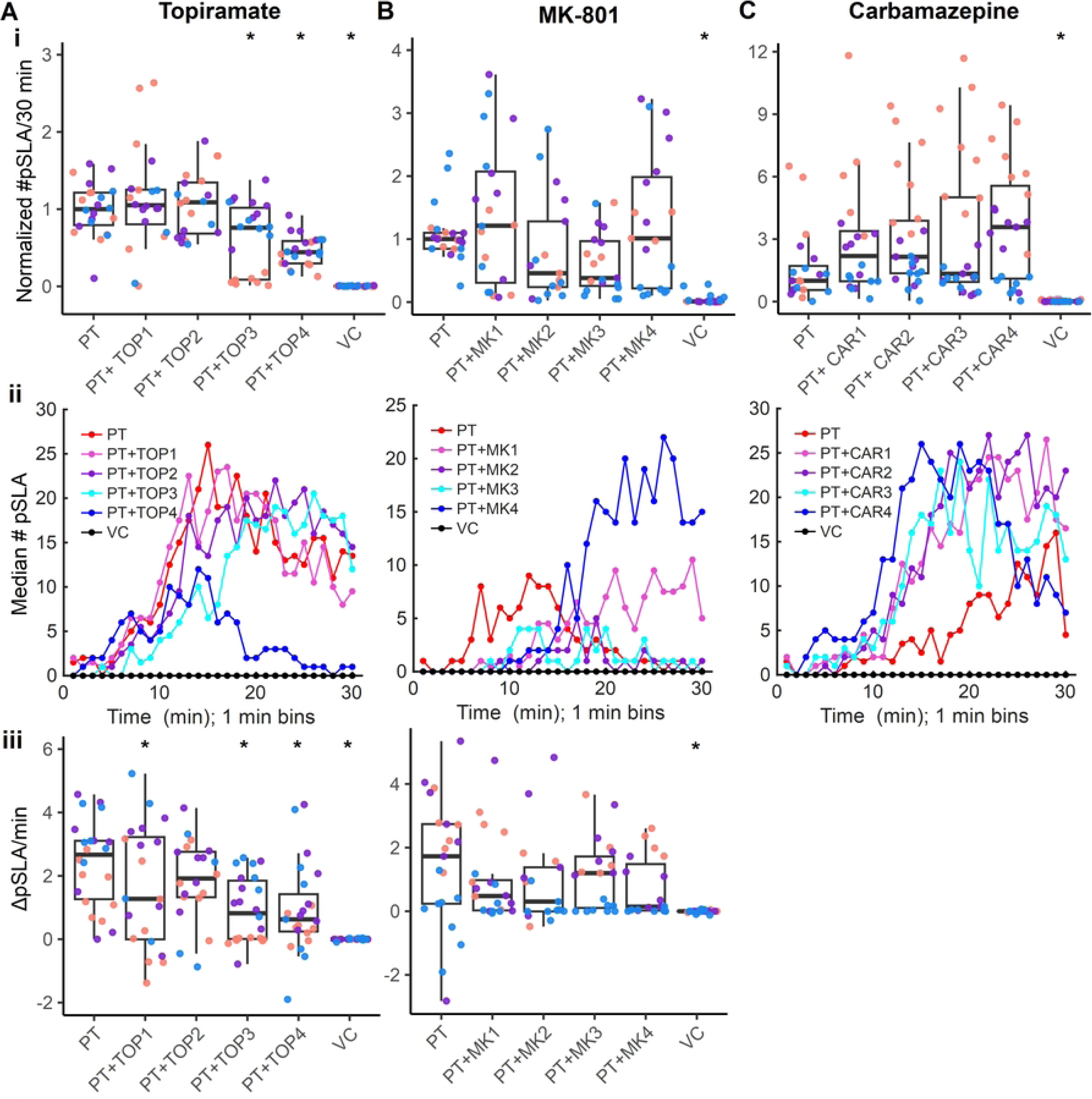
ASMs modulate parathion pSLA onset in DJ planarians. A-C) Effects of co-exposure of parathion (PT) and (A) topiramate (TOP) : 100, 316, 1000, 3160 µM; (B) MK-801 (MK): 1, 3.16, 10, 31.6 µM; or (C) carbamazepine (CAR): 10, 31.6, 100, 316 µM) were characterized by quantifying the normalized number of pSLA events after 30 minutes (i), visualizing pSLA dynamics over time (ii) and the change in pSLA per minute during the linear portion of the pSLA response (iii). Asterisk (*) indicates statistical significance (p<0.05) using a Kruskal Wallis omnibus test, followed by a post hoc one-sided Dunn’s test comparing each condition to the inducer alone with p-value correction using the BH method. VC: Vehicle control. Color-coding of individual data points in the boxplots indicates 3 independent experiments.

Co-exposure of parathion with up to 10 µM MK-801 caused a slight decrease in pSLA, which qualitatively appeared to be due to a delay in pSLA induction. However, due to the high variability observed across runs, none of the effects on the total number of pSLA or ΔpSLA/min were significantly different from parathion alone (Fig 3B). No significant effects on translational activity were observed with co-exposure to any MK-801 concentration (S5B Fig).

Carbamazepine was found to significantly increase the total number of pSLA and to cause pSLA onset earlier than parathion alone (Fig 3C). Because of this enhancing effect of carbamazepine on pSLA, we did not evaluate ΔpSLA/min. No effects on translational activity were observed with parathion and carbamazepine co-exposure (S5C Fig).

As was observed with midazolam, co-exposure of parathion with these three ASMs had different effects on parathion-induced pSLA in GD planarians than observed for DJ planarians (S6 Fig). Directly opposing results were found for topiramate and carbamazepine: topiramate decreased pSLA in DJ planarians but increased pSLA in GD planarians; carbamazepine increased pSLA in DJ planarians but decreased pSLA in GD planarians (S6 Fig). Decreases in parathion-induced pSLA with carbamazepine co-exposure were not correlated with decreased translational activity in GD planarians, suggesting a successful rescue. Interestingly, in GD planarians, carbamazepine seemed to show an opposite concentration-response as lower doses better attenuated pSLA, while the highest tested concentration of 100 µM returned to similar pSLA levels as in parathion alone, suggesting that different mechanisms may play a role at different concentrations. MK-801 was the only therapeutic to show similar effects on parathion-induced pSLA in the two species as 10 µM MK-801 lowered pSLA in DJ planarians and significantly reduced pSLA in GD planarians exposed to parathion.

### ASMs cause different effects when co-administered with other seizurogenic compounds: PTZ and NMDA

Given the variable results with parathion-induced pSLA and the stark differences between the two planarian species, we asked whether co-exposure of these ASMs with other seizurogenic chemicals was more consistent. PTZ is a commonly used seizurogenic compound, often used to create rodent and zebrafish seizure models [57–62], and we previously demonstrated that it induces pSLA in both DJ and GD planarians [46]. Therefore, we also evaluated the effects of the four ASMs on pSLA induced by exposure to 15 mM PTZ using co-exposure for 20 minutes. In DJ planarians, midazolam, topiramate and MK-801 had no effect on pSLA (Fig 4) or translational activity (S7 Fig). While carbamazepine also did not reduce pSLA in response to PTZ, there was a moderate increase in pSLA with higher concentrations (S8 Fig).

**Fig 4.**
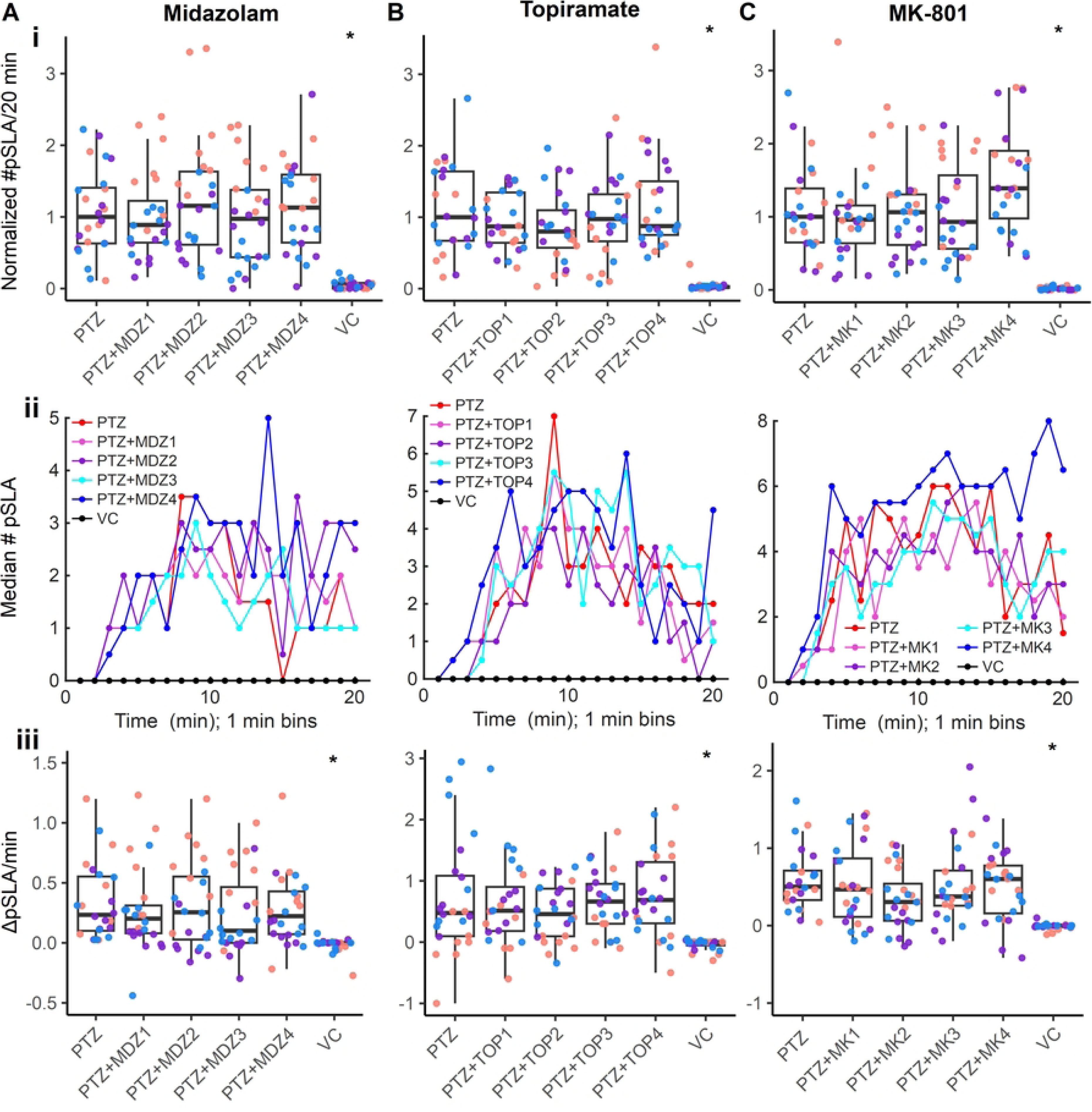
Midazolam, topiramate and MK-801 have no effect on PTZ-induced pSLA in DJ planarians. DJ planarians were co-exposed to 15 mM PTZ and either (A) midazolam: 0.1, 0.316, 1, and 3.16 µM; (B) topiramate (TOP): 100, 316, 1000, 3160 µM; or (C) MK-801 (MK): 1, 3.16, 10, 31.6 µM for 20 minutes. VC: Vehicle control. Effects on pSLA induction and dynamics were quantified as i) the total number of pSLA over the entire recording, normalized by the median of in-plate PTZ controls, ii) plots of median number of pSLA recorded each minute, and iii) ΔpSLA/min during the linearly increasing time period, determined by the PTZ- only controls from each plate. Asterisk (*) indicates statistical significance (p<0.05) using a Kruskal Wallis omnibus test, followed by a post hoc one-sided Dunn’s test comparing each condition to PTZ alone with p-value correction using the BH method. Color-coding of individual data points in the boxplots indicates 3 independent experiments.

In contrast, in GD planarians, co-exposure with 316, 1000, or 3160 µM topiramate significantly decreased the amount of PTZ-induced pSLA (S9Ai Fig). However, these effects were concomitant with a significant rapid decrease in translational activity (S9Aii Fig). GD planarians co-exposed to PTZ and topiramate were contracted into small ball-like shapes by the end of the 20-minute recording (S10 Fig), suggesting that topiramate worsened the effects of PTZ. MK-801 and carbamazepine showed no significant reduction in PTZ-induced pSLA, while co-exposure with 316 µM MK-801 increased pSLA (S9B-C Fig).

Topiramate, MK-801, and carbamazepine have previously been found to be effective against pSLA induced by NMDA in GD and *G. tigrina* [47,48]. Given the varied results with these ASMs for pSLA induced by parathion or PTZ, we tested whether we could recapitulate some of the previously published findings and could ameliorate pSLA in response to NMDA. Neither topiramate nor carbamazepine reduced pSLA induced by NMDA in either DJ or GD planarians; instead, topiramate and the highest concentration of carbamazepine showed increased pSLA in DJ planarians (S11 Fig).

## Discussion

### ASM efficacy at ameliorating pSLA is proconvulsant- and species-specific

Therapeutic effects that decreased or delayed pSLA were observed in co-exposures with parathion, with all four tested ASMs (midazolam, topiramate, MK-801, and carbamazepine) showing effectiveness in one planarian species. This demonstrates that pSLA can be alleviated by common ASMs. Moreover, to the best of our knowledge, this was the first time that the effectiveness of midazolam as a therapeutic has been demonstrated in planarians. However, the tested ASMs caused different results depending on the planarian species used (Fig 5). None of the tested ASM combinations with PTZ or NMDA were successful at reducing pSLA (Fig 5). While our statistical analysis only tested for decreased pSLA compared to the proconvulsant alone, we also observed qualitatively obvious increases in pSLA in some combinations, suggesting additive effects, though these effects were also species-specific (Fig 5).

**Fig 5.**
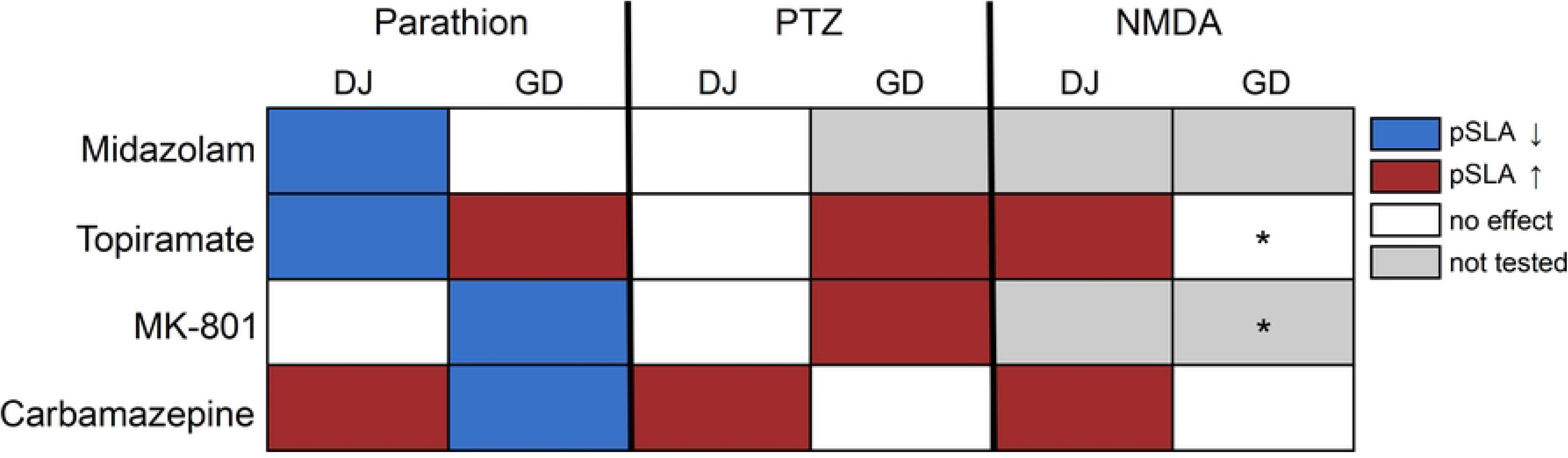
Overview of ASM effects across proconvulsants and planarian species. Therapeutic effects which decreased pSLA in the absence of toxicity are shown in blue. Aggravating effects that increased pSLA or caused overt toxicity are shown in red. Non-significant effects on pSLA are shown in white. Combinations which were not tested are indicated in gray. * indicates combination was tested previously by others in the literature [47].

Some of the ASMs tested here have previously been shown to reduce pSLA in certain conditions using manual scoring. Topiramate and MK-801, both tested at 0.1, 1, and 3 mM, have previously been shown to decrease NMDA-induced pSLA in GD planarians within 5 minutes [47]. We did not test MK-801 in combination with NMDA in GD planarians but topiramate did not reduce NMDA-induced pSLA under our experimental conditions. However, topiramate did reduce pSLA in response to parathion in DJ planarians, while MK-801 was successful in response to parathion in GD planarians. NMDA presented challenges for evaluating ASM effects in our experimental paradigm because it induced a rapid behavioral response, with planarians often crawling out of their wells within 5 minutes of NMDA exposure [46]. Therefore, we only analyzed effects within the first 5 minutes of recording and used additional plates to increase the sample size, excluding worms that crawled out of their wells during the 5-minute recording period. This crawl-out behavior was not mentioned as a confounding factor in the previously published results [47], likely because they used larger arenas (5.5 cm diameter petri dishes vs. 11 mm diameter wells) and mL versus µl volumes. In addition, we previously showed that NMDA solutions at these concentrations have a low pH that can independently induce pSLA [46]; thus, some of the behavioral effects of NMDA are likely driven by pH, which is not ameliorated by ASMs (S2 Table). Given these limitations, we did not evaluate all ASMs for effects on NMDA-induced pSLA.

In *G. tigrina* planarians, carbamazepine (1-100 µM) has been shown to reduce the number of pSLA events induced by NMDA, nicotine, and the GABA_A_ receptor antagonist picrotoxin in 5-minute recordings [48]. While neither species tested here showed attenuation of NMDA-induced pSLA by carbamazepine, carbamazepine reduced parathion-induced pSLA in GD planarians. In contrast, in DJ planarians, carbamazepine (particularly at 316 µM) increased pSLA when co-exposed with any of the proconvulsants (parathion, PTZ or NMDA). Thus, given the species-specific variability observed here, discrepancy with the previously published results could be due to species differences. While carbamazepine is a commonly used ASM for focal seizures in humans due to its inactivation of voltage-gated sodium channels [63], it often aggravates seizures in patients with generalized seizure types, such as absence seizures, by potentiating GABA_A_ receptors [64]. It remains to be determined whether the aggravating effects of carbamazepine seen with all proconvulsants in DJ planarians here is also due to effects on GABA_A_ receptors.

Similar to the inter-species variability observed for the ASMs here, we previously demonstrated that some chemicals differ in seizurogenic potential or potency in DJ and GD planarians [46]. Thus, there may be species differences in chemical uptake or receptor presence/identity that require further investigation. Overall, there was more variability in GD planarian behaviors compared to DJ planarians. This may be because GD planarians are commercially sourced and there can be behavioral and sensitivity differences between vendors and between batches from the same vendor, whereas the DJ planarians were from an established lab culture. Additionally, the GD pSLA response is relatively weaker, both in terms of the quantity of pSLA events observed and in the amount of pixel changes observed during a pSLA event [46]. Notably, previous pSLA literature relied on manual, qualitative definitions of pSLA that were broad (“asynchronous paroxysms”) and included C- and corkscrew shapes and other abnormal behaviors [47,48]. It is possible that not all of these behaviors would be detected as pSLA in our automated analysis which requires a certain threshold of movement. This may partially explain why previous studies were more successful at mitigating pSLA in GD planarians. Because of these multiple challenges, which make it more difficult to parse effects with the ASMs in GD planarians, we propose that DJ planarians are the preferred species for proconvulsant and ASM screening.

An important caveat to consider when interpreting the experiments presented here is that pSLA is a convulsive behavior that is used as a proxy for seizures [46]. Neuronal activity measurements during pSLA have not yet been conducted and it remains to be verified if pSLA is correlated with neuronal seizure activity and whether all seizure activity in planarians causes convulsions. For example, in developing zebrafish exposed to PTZ and other GABA_A_ receptor antagonists, it has been shown that ASM-mediated effects on locomotion are not always correlated with changes in electrographic neuronal activity as some ASMs decreased seizures without effects on behaviors and some ASMs showed behavioral effects in the absence of effects on seizures [23,65]. Similarly, in medicinal leech, co-exposure with the ASM phenytoin reduced neuronal hyperactivity *in vitro*, but did not decrease PTZ-induced hyperactivity [66]. It is possible that in our experiments the ASMs may have successfully induced neuronal changes that either did not manifest as behavioral changes or were unable to be detected with our analysis.

### Molecular targets and mechanisms in planarians

In humans, the proconvulsants and ASMs tested here span various mechanisms from specific effects on excitatory neurotransmission via glutamate or acetylcholine or inhibitory neurotransmission via GABA to broad effects on neuronal firing. The extent that these same mechanisms play a role in planarians is unclear. However, the human mechanisms can still provide a starting point for interpreting the different effects observed here.

Modulation of seizure activity is often achieved by regulating the balance of excitatory neurotransmission via glutamate and inhibitory neurotransmission via GABA. Thus, GABA_A_ receptor antagonists such as PTZ are often used to create seizure animal models [58,65]. Planarians have GABAergic neurons which are important for phototactic behavior [67], but the receptors have not yet been identified. It has previously been shown in GD planarians that exposure to 10 µM midazolam induces withdrawal-like behaviors, as locomotor activity decreases when midazolam is removed. Co-exposure with the GABA_A_ receptor antagonist flumazenil rescues this locomotor decrease [50], suggesting that midazolam acts on planarian GABA_A_ receptors. Given the presumed target conservation and the observed effectiveness of midazolam to decrease parathion-induced pSLA, we expected midazolam to decrease pSLA induced by PTZ in DJ planarians (this combination was not tested in GD planarians), as in vertebrates [65,68], but no effect was observed. In mammals, PTZ binds to the picrotoxin site of GABA_A_ receptors within the channel pore, stabilizing the closed channel conformation [69], while benzodiazepines like midazolam bind allosterically to the extracellular benzodiazepine site to potentiate channel opening [70,71]. Despite acting on separate sites, PTZ may induce conformational changes that inhibit benzodiazepine binding, but this has only been demonstrated at very high concentrations in rat brain synaptosomal membranes that may not be physiologically relevant to its convulsive action [72]. It is possible that differences in the planarian GABA_A_ receptor enable this interaction at lower, more behaviorally relevant concentrations. Midazolam and other benzodiazepines have been shown to be effective at mitigating seizures and hyperactive behavioral effects due to PTZ and other GABA_A_ receptor antagonists, picrotoxin and tetramethylenedisulfotetramine, in developing zebrafish larvae, but showed differential potency depending on which GABA_A_ receptor antagonist was used [65]. Effective concentrations needed to decrease locomotor effects induced by PTX were several-fold higher than for PTZ and tetramethylenedisulfotetramine. It is possible that DJ planarians have a narrow therapeutic window that we failed to cover in our experimental design.

In contrast to modulation of inhibitory neurotransmission via GABAergic signalling, seizures can also be induced by increased excitotoxicity via glutamatergic and cholinergic signalling. NMDA induces seizures via activation of NMDA glutamate receptors. Two putative genes encoding glutamate receptors in DJ planarians have been identified and their gene expression characterized [73], but their physiological functions and pharmacological sensitivities remain to be determined. The NMDA receptor antagonist MK-801 was previously shown to decrease NMDA-induced pSLA in GD [47] and *G. tigrina* [74]. Topiramate is a broad-spectrum drug believed to impart its anti-seizurogenic properties through multiple mechanisms, including inhibition of voltage-gated sodium and calcium channels, potentiation of GABA_A_ receptors, and antagonism of glutamate receptors [75]. In GD planarians, topiramate was previously found to decrease pSLA in response to NMDA, AMPA [47], and L-glutamate [49], suggesting that its functions on glutamate receptors may be conserved in planarians. Taken together, these data would suggest that the role of glutamate receptors in pSLA is functionally similar to that in mammalian seizures.

Parathion and other OPs induce seizures by inhibiting AChE, leading to accumulation of acetylcholine and subsequent overstimulation of central muscarinic acetylcholine receptors. We have previously shown that parathion is a potent inhibitor of DJ cholinesterase activity [28,36]. Two cholinesterases have been identified and characterized in DJ planarians and have been shown to be inhibited by OPs similarly to mammalian AChE [36,76]. Importantly, DJ planarians have also been shown to contain the major machinery necessary to metabolize OPs [45]. Previous work using atropine and tubocurarine has shown that both muscarinic and muscle nicotinic receptors contribute to the behavioral response to elevated acetylcholine in DJ [77] and that pSLA can be induced by the muscarinic receptor agonist pilocarpine in DJ and GD planarians [46,78]. However, muscarinic and nicotinic receptors remain to be molecularly characterized in both DJ and GD planarians. In mammals, OP-induced cholinergic toxicity is thought to trigger secondary neuronal changes that further promote seizure activity including increases in synaptic glutamate levels, overactivation of NMDA receptors, increased calcium influx, and lack of GABAergic inhibition [79]. This time-dependent shift from cholinergic- to glutamatergic-driven processes is why benzodiazepines like midazolam have a short time window for effectiveness against OP-induced seizures [1,8].

In summary, although planarians possess the same neurotransmitter systems and putative receptors implicated in vertebrate seizure activity, the pharmacological properties and physiological functions of many of these receptors remain poorly understood. Consequently, it is not yet clear whether ASMs that act on mammalian targets produce comparable molecular effects in planarians and will require mechanistic studies to unravel.

### Limitations and opportunities

The findings presented here demonstrate that some ASMs alleviate pSLA in response to parathion treatment, depending on the species, but could not alleviate pSLA in response to common seizurogenic compounds, PTZ and NMDA. Some of the results from this analysis conflict with published literature, emphasizing the need for replication studies to ensure findings are robust and generalizable. Importantly, all previous evaluations of MK-801, topiramate, and carbamazepine in planarians relied on manual counting of pSLA events, which were qualitatively characterized, over the course of a 5-minute exposure. The automated image analysis pipeline used here thus provides a robust, quantitative metric of pSLA quantity and temporal dynamics that was not captured by previous analyses and can be more consistently compared across studies. While we evaluated most proconvulsant + ASM combinations for pSLA quantity and dynamics, and translational activity, this pipeline could be streamlined into a tiered approach in the future and certain improvements can be made to further augment its value.

A major challenge associated with these quantitative analyses was the high variability observed across experiments. As can be seen from comparing the pSLA over time curves across experiments and from the across individual spreads in ΔpSLA/min values, the timing and magnitude of pSLA induction with parathion could vary considerably from plate to plate. We tried to minimize variability from factors such as time of day, fasting state, and worm size, but did not systemically study their contribution. To try to control for this variability and correct for any chemical batch effects, all statistically evaluated responses (total number of pSLA, ΔpSLA/min, translational AUC) were normalized by the responses of the proconvulsant-only controls in the same plate. Even with this normalization, some conditions (such as the parathion + MK-801 co-exposures) still showed large variability across worms, even within the same plate, decreasing statistical power and limiting our ability to detect effects with the ASMs. Better understanding and control of this worm-to-worm variability in pSLA induction will be needed if planarians are to be used to screen for novel ASMs.

While our analysis pipeline can quantify and compare pSLA quantity and rate of onset, it cannot differentiate different types of hyperactive behaviors because pSLA is defined as activity exceeding a threshold of pixel changes [46]. Thus, any behaviors above that threshold are indiscriminately detected as pSLA. Our motility index-based measure mimics approaches in other small model organisms that rely on a general metric of overall activity, such as total distance moved [65] or the sum of pixel differences between time intervals [23] (equivalent to motility index used to define pSLA here [46]), to approximate convulsive hyperactivity and thereby interpret loss of hyperactivity as a successful rescue of seizurogenic activity. However, there are examples in the literature showing that these generic measures can fail to detect behavioral changes that have similar levels of motility. For example, in the medicinal leech, pre-treatment with the ASM phenytoin could not rescue PTZ-induced hyperactivity but reduced the amount of abnormal twisting or tumbling behaviors and reduced neuronal hyperactivity *in vitro* [66]. Thus, it is possible that in our experiments some ASMs may have altered neuronal signalling in a manner that our behavioral analysis could not detect.

On the flip side, loss of hyperactivity is not always associated with antiseizurogenic effects. In developing zebrafish, 18-hour pre-treatment to the ASM zonisamide decreased PTZ-induced hyperactivity but worsened seizures measured by electroencephalography [23]. The authors speculate that this is because zonisamide caused prolonged late-stage seizures wherein the larvae move less due to loss of posture and balance, which may be similar to status epilepticus in rodents. In our study, we quantified both hyperactive events (pSLA) and translational activity to be able to better interpret whether decreased pSLA was indicative of a rescue or aggravated effects. However, both wild-type gliding behavior and pSLA have high translational activity; thus, making it difficult to interpret whether decreased translational activity resulting from ASM co-exposure is a result of loss of pSLA, as with parathion + midazolam in DJ planarians, or due to toxicity, as in PTZ + topiramate in GD planarians, without qualitative analysis of the recordings. Therefore, displacement alone is likely insufficient for detecting and characterizing ASM activity. Additional metrics, such as those evaluating body shape changes or directed motion, will be needed to allow for automated, quantitative ways to distinguish these various classes of effects. For example, shape mode analysis has been used to characterize and differentiate different shape classes in nematodes [80] and planarians [81] and distinguish behavioral responses. These could be combined with machine learning approaches, such as recently developed to detect seizure activity in zebrafish [82]. In addition, stimulus-evoked behaviors such as light-dark transitions, as used in developing zebrafish [83] and additional behavioral readouts, such as thigmotaxis or phototaxis, are worth exploring.

## Conclusions

A central finding of this study was that the effects of ASMs on pSLA depended on both the proconvulsant used to induce the behavior and the planarian species. No single ASM consistently attenuated pSLA across all conditions, and some treatments had opposite effects depending on the proconvulsant or species. This variability suggests that the effectiveness of an ASM may depend on the mechanism used to induce pSLA and how structurally similar the molecular targets in the planarian species studied are to the human targets. Systematic characterization of planarian receptors and signalling pathways, combined with testing compounds with defined mechanisms of action, will be important for determining which behavioral phenotypes are representative of a given mechanism of action. This mechanistic foundation will be essential for determining the potential of planarians as a model for identifying novel ASMs.

## Acknowledgements

The authors thank Matthew Shiley for making the Fresnel lenses used in the imaging set-up, Hannah Poon for the planarian schematic in Fig 1, Dr. Hilary McCarren, Antonia Santoro, Julia Morgan, and Dr. Rudy J. Richardson for discussions, Dahlia Bedward and Sophia Kral for help with planarian care.

## Supporting information captions

**S1 Fig. Dose response for antiseizure medications (ASMs) in DJ.** DJ planarians were exposed to either midazolam, topiramate, MK-801, or carbamazepine for 20-30 minutes and A) the total number of pSLA events over the entire recording and B) translational activity normalized by worm area were quantified. VC: Vehicle control. All concentrations are in µM. Color-coding in A indicates 3 independent experiments. Asterisk (*) indicates statistical significance (p<0.05) determined from pairwise contrasts (with a Benjamini & Hochberg p-value correction) of the estimated marginal means of a negative binomial generalized linear model of concentration versus number of pSLA events. Translational activity plots show the median results for each condition across n=24 planarians, from 3 independent experiments.

**S2 Fig. Dose response for antiseizure medications (ASMs) in GD.** GD planarians were exposed to either midazolam, topiramate, MK-801, or carbamazepine for 30 minutes and A) the total number of pSLA events over the entire recording and B) translational activity normalized by worm area were quantified. VC: Vehicle control. All concentrations are in µM. Color-coding in A indicates 3 independent experiments. Asterisk (*) indicates statistical significance (p<0.05) determined from pairwise contrasts (with a Benjamini & Hochberg p-value correction) of the estimated marginal means of a negative binomial generalized linear model of concentration versus number of pSLA events. Translational activity plots show the median results for each condition across n=24 planarians, from 3 independent experiments.

**S3 Fig. Midazolam delays parathion-induced pSLA in DJ planarians.** A) DJ planarians were co-exposed to 75 µM parathion (PT) and varying concentrations of MDZ (MDZ1-4 correspond to 0.1, 0.316, 1 and 3.16 µM, respectively) and total number of pSLA events, normalized by in-plate parathion-only controls, over the entire 90-minute recording were counted. Asterisk (*) indicates statistical significance (p<0.05) using a Kruskal Wallis omnibus test, followed by a post hoc two-sided Dunn’s test comparing each condition to parathion alone with p-value correction using the BH method. Color-coding of individual data points indicates 2 independent experiments. VC: Vehicle control. B) Dynamics of pSLA induction were visualized by plotting the median number of pSLA for each minute, showing that all concentrations, except for MDZ1, eventually induce pSLA. C) Translational activity (from 2 independent experiments) was quantified from minimum intensity projections and normalized by each worm’s area.

**S4 Fig. Midazolam has no effect on parathion-induced pSLA in GD planarians.** A) GD planarians were co-exposed to 31.6 µM parathion (PT) and varying concentrations of MDZ (MDZ1-4 correspond to 0.1, 0.316, 1 and 3.16 µM, respectively). Total number of pSLA events, normalized by in-plate parathion-only controls, over the entire 30-minute recording were counted. B-C) Translational activity was quantified from minimum intensity projections and normalized by each worm’s area. Plots of translational activity over time (median of all worms) show all PT-containing exposures quickly decrease their translational motion (B) and are quantified by comparing the area under the curve (AUC) (C). Asterisk (*) indicates statistical significance (p<0.05) using a Kruskal Wallis omnibus test, followed by a post hoc two-sided Dunn’s test comparing each condition to the inducer alone with p-value correction using the BH method. Color-coding of individual data points in A and C indicates 3 independent experiments.

**S5 Fig. Effects of topiramate, MK-801 and carbamazepine on translational activity when co-exposed with parathion in DJ planarians.** A-C) Effects of co-exposure of parathion (PT) and (A) topiramate (TOP): 100, 316, 1000, 3160 µM; (B) MK-801 (MK): 1, 3.16, 10, 31.6 µM; or (C) carbamazepine (CAR): 10, 31.6, 100, 316 µM) on translational activity. VC: Vehicle control. Plots of median translational activity over time (top) and their quantification by comparing the area under the curve (AUC) (bottom) are shown. Asterisk (*) indicates statistical significance (p<0.05) using a Kruskal Wallis omnibus test, followed by a post hoc two-sided Dunn’s test comparing each condition to the inducer alone with p-value correction using the BH method. Color-coding of individual data points indicates 3 independent experiments. D) Representative images of VC, PT only and PT+TOP4 at the end of the 30-minute recording to show abnormal morphology observed with PT+TOP4 concomitant with decreased translational activity. Arrowhead indicates pharynx that was ejected from the body. Scale: 1 mm.

**S6 Fig. Topiramate, MK-801 and carbamazepine modulate parathion-induced pSLA in different ways in GD planarians.** A-C) Effects of co-exposure of 31.6 µM parathion (PT) and (A) topiramate (TOP): 100, 316, 1000, 3160 µM; (B) MK-801 (MK): 1, 3.16, 10, 31.6 µM; or (C) carbamazepine (CAR): 3.16, 10, 31.6, 100 µM) in GD planarians were characterized by (i) quantifying the normalized number of pSLA events after 30 minutes. (ii-iii) Translational activity was quantified from minimum intensity projections and normalized by each worm’s area. Plots of median translational activity over time (ii) and their quantification by comparing the area under the curve (AUC) (iii) are shown. Asterisk (*) indicates statistical significance (p<0.05) using a Kruskal Wallis omnibus test, followed by a post hoc one-sided Dunn’s test comparing each condition to the inducer alone with p-value correction using the BH method. Color-coding of individual data points in the boxplots indicates 3 independent experiments.

**S7 Fig. Midazolam, topiramate and MK-801 have no effect on PTZ translational activity in DJ planarians.** DJ planarians were co-exposed to 15 mM PTZ and either (A) midazolam: 0.1, 0.316, 1, 3.16 µM; (B) topiramate (TOP): 100, 316, 1000, 3160 µM; or (C) MK-801 (MK): 1, 3.16, 10, 31.6 µM for 20 minutes. Effects on translational activity were characterized via plots of median translational activity over time (i) and their quantification by comparing the area under the curve (AUC) (ii) None of the conditions were significantly different (p<0.05) using a Kruskal Wallis omnibus test, followed by a post hoc one-sided Dunn’s test comparing each condition to PTZ alone with p-value correction using the BH method. Color-coding of individual data points in the boxplots indicates 3 independent experiments.

**S8 Fig. Carbamazepine increases pSLA in response to PTZ in DJ planarians.** DJ planarians were co-exposed to 15 mM PTZ and carbamazepine (CAR): 10, 31.6, 100, 316 µM for 20 minutes. A) The total number of pSLA events, normalized by in-plate PTZ-only controls, were counted. B-C) Dynamics of pSLA induction were visualized by (B) plotting the median number of pSLA for each minute and (C) calculating the ΔpSLA/min during the linearly increasing time period, determined by the PTZ-only controls from each plate. D-E) Translational activity was quantified from minimum intensity projections and normalized by each worm’s area. Plots of translational activity over time (D) were quantified by comparing the area under the curve (AUC) (E). Asterisk (*) indicates statistical significance (p<0.05) using a Kruskal Wallis omnibus test, followed by a post hoc one-sided Dunn’s test comparing each condition to the inducer alone with p-value correction using the BH method. Color-coding of individual data points in the boxplots indicates 3 independent experiments.

**S9 Fig. Effects of topiramate, MK-801, and carbamazepine on PTZ pSLA and translational activity in GD planarians.** GD planarians were co-exposed to 15 mM PTZ and either (A) topiramate (TOP): 100, 316, 1000, 3160 µM; or (B) MK-801 (MK): 1, 3.16, 10, 31.6 µM; or (C) carbamazepine (CAR): 3.16, 10, 31.6, 100 µM for 20 minutes. i) Effects on pSLA induction were quantified as the total number of pSLA over the entire recording, normalized by the median of in-plate PTZ controls. Effects on translational activity were characterized via ii) plots of median translational activity over time and iii) their quantification by comparing the area under the curve (AUC) (bottom). Asterisk (*) indicates statistical significance (p<0.05) using a Kruskal Wallis omnibus test, followed by a post hoc one-sided Dunn’s test comparing each condition to PTZ alone with p-value correction using the BH method. Color-coding of individual data points in the boxplots in indicates 3 independent experiments.

**S10 Fig. Topiramate causes paralysis in PTZ-treated GD planarians.** Representative images of GD planarians exposed to vehicle control (VC), 15 mM PTZ, or PTZ and 100 µM topiramate (PTZ+TOP1). Images are from the last 15-20 minutes of exposure. PTZ+TOP exposed planarians are highly contracted into small balls with almost no movement.

**S11 Fig. Topiramate and carbamazepine do not rescue NMDA-induced pSLA in DJ or GD planarians.** A-B) DJ planarians were co-exposed to 15 mM NMDA and either (A) topiramate (TOP): 100, 316, 1000, 3160 µM; or (B) carbamazepine (CAR): 10, 31.6, 100, 316 µM for 5 minutes and the total amount of pSLA, normalized by in-plate NMDA controls, were compared. C-D) GD planarians were co-exposed to 3 mM NMDA and either (A) topiramate (TOP): 100, 316, 1000, 3160 µM; or (B) carbamazepine (CAR): 3.16, 10, 31.6, 100 µM for 5 minutes and the total amount of pSLA, normalized by in-plate NMDA controls, were compared. Asterisk (*) indicates statistical significance (p<0.05) using a Kruskal Wallis omnibus test, followed by a post hoc one-sided Dunn’s test comparing each condition to the inducer alone with p-value correction using the BH method. Color-coding of individual data points in the boxplots indicates 3-4 independent experiments.

**S1 Table. pH of highest test concentrations of each chemical.**

**S2 Table. pH of highest test concentrations used in co-exposures.**

**S3 Table. P-values and sample sizes.**

**S1 Movie. Loss of translational motion in DJ planarians exposed to parathion and midazolam is not due to toxicity.** Example movies of planarians exposed to the vehicle control, parathion (PT) or PT and 0.1 µM midazolam (PT + MDZ) taken from the last 5 minutes of the 30-minute recording. Although PT+MDZ worms have decreased translational motion, they do not exhibit obvious morphological abnormalities to suggest toxicity. Videos are at 2X speed. The diameter of each well is 11 mm.

